# Contrasting temporal patterns and associations in *Hyalomma marginatum* microbial communities: key insights for the development of novel tick and tick-borne diseases control tools

**DOI:** 10.1101/2025.05.21.655249

**Authors:** Joly-Kukla Charlotte, Manzanilla Vincent, Duhayon Maxime, Bru David, Mélanie Jeanneau, Moutailler Sara, Pollet Thomas

## Abstract

**Background:** *Hyalomma marginatum* is an invasive tick species in southern mainland France that can carry several pathogens of human and animal interest. Because the tick microbiota represents a major factor that can potentially modulate the pathogen acquisition and transmission and might become a new control tool against ticks and tick-borne diseases, it is more than essential to identify the composition of the *H. marginatum* microbiota, its temporal dynamics and interactions (statistical association) between members of the tick microbiota.

**Methods:** From monthly tick samplings performed in the same site between February to September 2022, 281 adult ticks *H. marginatum* were collected from horses. The microbiota composition was characterised using a high throughput sequencing approach. Different statistical analyses allowed us to assess the influence of several factors (month, season, tick sex) on the *H. marginatum* microbial communities and reveal potential interactions among members of these communities.

**Results:** Apart of known obligate endosymbionts *Francisella* and *Midichloria*, and the hypothesised facultative endosymbiont (*Rickettsia*) that dominated the microbiota of *H. marginatum*, we detected *Staphylococcus*, *Corynebacterium*, *Williamsia* and *Mycobacterium*, usually described as commensal and/or environmental bacteria. The microbiota composition and bacterial networks differed between males and females, with males being more diverse and composed of more environmental bacteria. We reported several temporal shifts for both males and females into the microbiota composition and bacterial networks. The temporal shifts observed for females were more chaotic in terms of movements among nodes, compared to the male microbial communities that exhibited a more organised and stable dynamics.

**Conclusions:** The influence of tick sex and time on the holobiont *H. marginatum* underlines the importance of the scale at which the study is conducted.

**Highlights:**

- Obligate endosymbionts dominate the *Hyalomma marginatum* microbiota
- Environnemental bacteria are abundant in *Hyalomma marginatum* microbiota
- Both the bacterial composition and associations differed according to the tick sex
- Both the bacterial composition and associations were marked by many temporal shifts for both males and females
- Keystone taxa in tick microbiota were highly variable from one month to another

## Background

Ticks transmit Tick-Borne Pathogens (TBP) affecting human and animal health. These pathogens coexist with non-pathogenic microbial communities associated with ticks, collectively known as the microbiota encompassing symbionts, commensals and environmental microbes (Bonnet *et al*. 2017; Vayssier-Taussat *et al*. 2014). The bacterial component that has been particularly studied in ticks in recent years plays an important role in the tick’s physiology, nutrition, development and reproduction (Bonnet *et al*. 2017; Bonnet et Pollet 2021; Duron *et al*. 2018; Narasimhan et Fikrig 2015). Ticks typically carry primary/obligate symbionts that are vital for the tick’s survival. In *H. marginatum* for example, *Midichloria* and *Francisella*-LE are two primary symbionts that provide B vitamins absent from the blood meal but vital for the tick’s survival (Buysse *et al*. 2021). Ticks also harbour secondary/facultative symbionts that are beneficial but not essential to their survival (Bonnet *et al*. 2017; Su, Zhou, et Zhang 2013). Their functions in ticks are still to be elucidated but their roles in the defence against abiotic or biotic stresses are hypothesised, as this is the case for *Spiroplasma ixodetis* that might be involved in the defence of *I. ricinus* ticks against parasitoid wasps (Lejal *et al*. 2021) and in the defence against pathogenic *Rickettsia* (Aivelo, Norberg, et Tschirren 2019). In addition to symbionts, other microbes can be acquired by ticks through the environment (*i.e.* the host skin, vegetation, soil), and can colonise the tick midgut (Bonnet *et al*. 2017; Ross *et al*. 2018; Zolnik *et al*. 2018). Beyond their contribution to tick physiology and bio-ecology, microbiota members potentially interact with TBP as hypothesised for the secondary symbiont *Rickettsia bellii* symbiont and the TBP *A. marginale* in *D. andersoni* (Gall *et al*. 2016). Similarly, interaction among environmental bacteria including *Bacillus*, *Pseudomonas* and *Enterobacteriaceae* were also found to be correlated to pathogens such as *B. burgdorferi s.l.* in *I. scapularis* (Ross *et al*. 2018). These potential interactions can affect the vector competence of various tick species, by facilitating or impairing the pathogen acquisition, maintenance or transmission (Pollet *et al*. 2020; Wu-Chuang *et al*. 2021). These advances paved the way to emerging tools for tick and tick-borne pathogens control strategies (Mateos-Hernández *et al*. 2021; Wu-Chuang *et al*. 2022a).

The tick holobiont, as referring to the biological and evolutionary entity composed of the tick and its microbial communities, is a dynamic entity that can vary along with tick-associated factors including the tick species, population, life stage, tick sex, the engorgement status and the anatomical location (Pollet *et al*. 2020). Space and time scales are also crucial to consider since the tick holobiont is likely to vary depending on where and when the ticks were collected (Pollet *et al*. 2020). By affecting tick’s bio-ecology (activity, metabolism), the spatial scale and temporality also influence the composition of the microbiota. Some studies have indeed identified substantial temporal variations into the microbiota composition in *Hyalomma dromedarii* from March 2019 to February 2020 in United Arab Emirates (Perveen *et al*. 2022), *Rhipicephalus sanguineus s.l.* and *R. turanicus* from March to July 2009 in Israel (Lalzar *et al*. 2012) and *Ixodes ricinus* between April 2014 and May 2017 in France (Lejal *et al*. 2021).

*Hyalomma marginatum* is an invasive tick species that recently became established in southern mainland France in 2015, in areas where the climate has become more suitable for this tick species, probably due to climate change (Stachurski et Vial 2018; Vial *et al*. 2016). It seems its expansion area keeps expanding in northern latitudes in France. This tick species is of particular interest for both human and animal health, indeed in previous studies, we identified several TBP carried by *H. marginatum* in southern mainland France between 2016 and 2024 (Bernard *et al*. 2024a; Joly-Kukla *et al*. 2024a), especially the Crimean-Congo fever haemorrhagic virus that was detected in *H. marginatum* ticks for the first time in France (vector and reservoir) (Bernard *et al*. 2024b). We also demonstrated that several of these TBP were highly affected by the temporal scale, especially for *R. aeschlimannii*, that was more prevalent during summer compared to spring and winter (Joly-Kukla *et al*. 2024b). Determining how the temporal patterns and other tick-related factors affect *H. marginatum* microbiota and its interactions (statistical association) increase our knowledge on tick-microbiota-pathogen interactions and make our contribution to the development of potential new control strategies of ticks and TBP. In this perspective, 281 *H. marginatum*, males and females, were collected at monthly intervals in 2022 between February and September in the same site next to Montpellier (South of mainland France). We then assessed the *H. marginatum* microbiota and its temporal dynamics using a high throughput sequencing approach. Using network analyses, we identified direct statistical interactions among members of the microbiota, including pathogenic genera, and assessed the temporal variability of these interactions. We inferred a static network that contained persistent and recurring interactions throughout the months for both males and females. In addition, we inferred monthly subnetworks to investigate the dynamic of microbial interactions.

## Methods

### Tick collection

Ticks were monthly collected from February to September 2022 at a site called Mas de la Lauze, in Pompignan, located in the Gard department in southern mainland France. All ticks were directly removed from horses. More details about the sampling sites and the tick collection are available in Joly-Kukla *et al*., 2024b. The sampling dates and the number of *H. marginatum* that were analysed were as follows: February 24th (n=21 ticks), March 11th (n=31), March 17th (n=24), March 31th (n=34), April 28th (n=34), May 25th (n=34), June 23th (n=31), July 21th (n=33), August 19th (n=28) and September 16th (n=11), giving a total of 281 *H. marginatum*.

### DNA extraction

Ticks were crushed individually and nucleic acids were extracted in a BSL3 facility. As previously described in Joly-Kukla *et al*., (2024b), the ticks were washed for 30 seconds in 1% hypochlorite solution that was diluted from a bottle containing 2.6% of active chlorine (Orapi Group, Saint-Vulbas, France) then rinsed three times for one minute in Milli-Q water to remove any environmental microbes present on the tick cuticle (Binetruy *et al*. 2019b). Ticks were then cut into pieces using a scalpel blade and crushed individually in a Precellys®24 Dual homogenizer (Bertin, France) at 5500 rpm for 40 sec, using three steel beads (2.8 mm, OZYME, Saint-Cyr-LLEcole, France) in 400 µL of DMEM (Dulbecco’s Modified Eagle Medium, Eurobio Scientific, Les Ulis, France) with 10% foetal calf serum. Total DNA and RNA was extracted using the NucleoMag VET extraction kit (Macherey-Nagel, Hoerdt, France) following the manufacturer’s instructions using the IDEALTM 96 extraction robot (Innovative Diagnostics, Grabels, France).

### 16S DNA Metabarcoding

DNA amplification was performed on the V4 region of the 16S rRNA gene using bar-coded 515F (16S-V4F: 5′-GTGCCAGCMG CCGCGGTAA-3′) and 806R (5′-GGACTACHVGGG TWTCTAATCC-3′) primers described by Galan *et al*. 2016; Lejal *et al*. 2021, producing a 251-bp amplicon. Different 8 bp indexes were added to primers allowing *in fine* the amplification and multiplexing of all samples. All the PCR amplifications were carried out using the Phusion® High-Fidelity PCR Master Mix kit (Thermo Fischer Scientific, Waltham, Massachusetts, USA). For each sample, 5 μL of DNA extract were amplified in a 50 μL final reaction volume. The following thermal cycling procedure was used: initial denaturation at 98 °C for 30 s, 35 cycles of denaturation at 98 °C for 10 s, annealing at 55 °C for 30 s, followed by extension at 72 °C for 30 s. The final extension was carried out at 72 °C for 10 min. Amplicons were checked on 1.5% agarose gels and normalised to ∼1ng/µL using the SequalPrepTM kit (Invitrogen Corp, Carlsbad, USA). Normalised amplicons were then pooled at equimolar concentrations and sent to the sequencing platform GenoToul (Toulouse, France). In addition to tick samples, 22 negative controls (washing, crushing, extraction, amplification controls) have been performed to distinguish tick microbial ASVs from contaminants (Lejal *et al*. 2020).

### Sequencing and data processing

The equimolar pool was sequenced by the GenoToul platform (Toulouse, France) using MiSeq Illumina 2 × 250 bp chemistry. The raw data were imported into the FROGS pipeline (Bernard *et al*. 2021; Escudié *et al*. 2018) for bioinformatic analyses (Escudié *et al*. 2018; Bernard *et al*. 2021). The demultiplexing was performed in order to trim barcodes and assign the sequences to the corresponding samples. After merging the reads using vsearch, sequences were trimmed to remove the primers using cutadapt. Resulting sequences were dereplicated, filtered by length (270-330pb) and pooled into ASVs (Amplicon Sequence Variant) using Swarm. Chimeras were removed with vsearch and a first filter was performed to keep ASVs representing at least 0.0005% of all sequences, as recommended by Bokulich *et al*., 2013. The taxonomic affiliation of ASVs was performed by blasting the seed ASV sequence with the 16S SILVA Pintail100 138.1 database. The taxonomic affiliation was performed as follows: kingdom, phylum, class, order, family and genus. ASVs corresponding to chloroplast sequences were removed.

To identify contaminant ASVs, a confidence interval corresponding to 1% ± 0.05% of the total number of sequences in the library was used. If the number of sequences of an ASV in the negative controls was higher than the upper threshold, it was considered as a contaminant ASV, removed from the dataset (Galan *et al*. 2016; Lejal *et al*. 2020). After the removal of negative control samples and contaminant ASV from the dataset, it was reuploaded into the FROGS pipeline in order to apply other ASV filters. More precisely, ASVs whose total number of sequences was lower than the total number of samples in the library were removed. In addition, ASVs detected in less than 15% of the samples were removed. In sum, the final dataset of 281 samples comprised 96 ASVs and 456, 577 sequences.

### Bacterial composition and diversity analysis

The analysis of composition and diversity was performed in R studio 4.3.3. The alpha diversity was determined using the richness (number of observed ASV), the Shannon index (ASV abundance distribution) and inverse Simpson index (inverse probability that two sequences in a given sample belong to the same ASV) using the *phyloseq* package (McMurdie et Holmes 2013). For the analysis of alpha diversity between males and females, generalised linear models (GLMs) with a Gamma distribution were used to assess the statistical differences in the Shannon and Inverse Simpson indices, while a linear model (LM) was used to analyse the richness (normal distribution) (package *lme4*, Bates *et al*., 2015). All models included both the tick sex and month as variables. The influence of months on alpha diversity was assessed for both males and females separately using LM and GLMs. Significance of variables and p-values were assessed using the “analysis of variance” procedure within the package *car* which performs a type III hypothesis (Fox et Weisberg 2018). The beta diversity was assessed with Bray-Curtis distances (fraction of the community that is specific to each of the samples compared), from the *phyloseq* package. The influence of the tick sex, month and season variables was determined with the *adonis2* (PERMANOVA) function of the *vegan* package (Oksanen *et al*. 2024).

The relative abundance of the genera was statistically analysed using GLMs with a Gamma distribution and using hurdle models for the process of zero counts and positive counts (package *pscl*, Jackman, 2024; Zeileis *et al*., 2008). We analysed the top 8 genera that represented more than 2.5% of monthly relative abundance in males, namely *Francisella*, *Midichloria*, *Rickettsia*, *Corynebacterium*, *Acinetobacter*, *Mycobacterium*, the multi-affiliated ASV of the *Bacilli* class and the multi-affiliated ASV of the *Moraxellaceae* family. For females, we analysed the top five genera above the 2.5% threshold namely *Francisella*, *Midichloria*, *Rickettsia*, *Corynebacterium* and the multi-affiliated ASV of the *Bacilli* class, but we also included into the analysis the other genera cited above for males, which were above this threshold for females, namely *Mycobacterium*, *Acinetobacter* and the multi-affiliated ASV of the *Moraxellaceae* family. A SIMPER (similarity percentage procedure) analysis based on the Bray-Curtis dissimilarities was performed to identify the contribution of ASVs to the difference of bacterial communities between males and females (simper, package *Stats Rbase* v4.1.3 Clarke, 1993).

As we performed the detection of tick-borne pathogens species using a high-throughput microfluidic based approach in our previous study (Joly-Kukla *et al*. 2024b) (‘microfluidic dataset’), and since we performed the microbiota analysis using the metabarcoding in the present study (‘ASV dataset’), we decided to integrate these two datasets using a multi-omics factor analysis (MOFA, package *MOFA2*). MOFA provides a general framework for the integration of multi-omic data sets (Argelaguet *et al*. 2018; Velten *et al*. 2022). We made the choice to not normalise the data, because it improved the variance explained by the factors (**Supplementary material**). The significance of the MOFA factors were assessed by the variance-explained estimates (per data modality) that resulted from the variance decomposition analysis.

The MOFA model was trained over 410 iterations with a convergence threshold of 0.1. The model’s performance indicated little correlation between the two modalities, (factor correlation of 0.06). The factors captured independent sources of variation from each modality. For the microfluidic dataset, the variance explained by the model was relatively low (3.64%) and the Tau value (0.25), which indicated that only a few features contributed significantly to the model. Given these observations, we decided not to exploit the microfluidic data results. In contrast, for the ASV data, the model was explaining 57.48% and the Tau value was much larger (5,874), indicating the model captured meaningful features.

### Network analysis

From the ASV dataset and based on the read counts, we modelled the relationship between ASVs under a Generalised Linear Model (GLM) assumption. Since our count data exhibited expected over-dispersion, we employed a multivariate Poisson log-normal (PLN) model, implemented in the *PLNmodels* package (Chiquet, Robin, et Mariadassou 2019). The offset was computed using Wrench normalisation. We considered data obtained for males and females as two separate datasets and generated male and female static networks that summarised significant interactions in time, by including month as a factor in the model. We included an intercept in the model to account for the baseline microbial abundance. We evaluated the performance of the PLN network model by comparing the fitted values with the observed values (**Figure S1**). To select the best network, we applied the Stability Approach to Regularization Selection (StARS), retaining only the strongest correlations with a stability set at 0.9 (supplemental material). The network was visualised using MetaNet. In addition to static networks, we generated monthly subnetworks for both males and females, to assess the temporal dynamics of bacterial interactions.

## Results

### Composition and diversity of *H. marginatum* microbiota

Considering the 281 samples, 96 ASVs comprising 456,577 sequences were identified. These ASVs belonged to five phyla, seven classes, 18 orders, 24 families and 27 genera. At the level of the genera, nine ASVs were multi-affiliated, three were multi-affiliated at the family level and one was multi-affiliated at the order level.

### Microbiota of male and female ticks

We first aimed to compare the microbiota of males and females ticks from all months. The 96 ASVs that were identified were common to males and females. The bacterial composition was firstly explored by identifying phyla and genera whose proportion of sequences was above the threshold of 0.5% of the total sequence number. For both male and female, three phyla were above the threshold: *Pseudomonadota*, *Actinomycetota* and *Bacillota*. The proportion of *Pseudomonadota* phylum was higher in females while the proportion of *Actinomycetota* and *Bacillota* were higher in males (**Figure 1A, 1B**). *Pseudomonadota*, *Actinomycetota* and *Bacillota* were encountered in almost all of the ticks whether males or females (**Figure 1C, 1D**). The phyla under the threshold were *Bacteroidota* and *Fusobacteriota*. These two phyla were both encountered more in males (in 74.5% and 52.9%) than females (60.2% and 46.1%).

**Figure 1:**
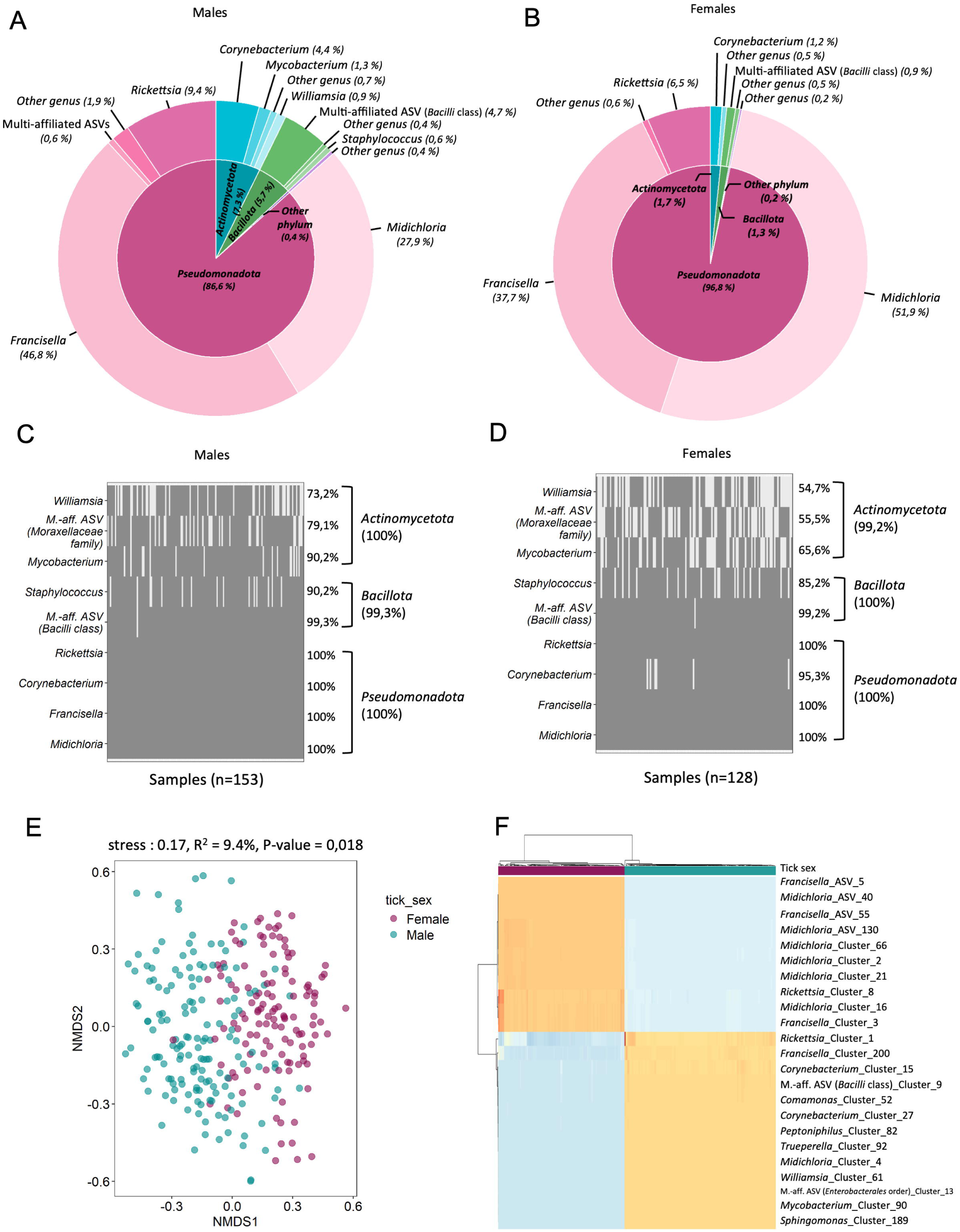
Influence of the tick sex on *H. marginatum* microbiota. Pie charts with phylum (inner circle) and genera (outer circle) of taxa whose number of sequences represent more than a threshold of 0.5% of total sequences in **A**. males or **B**. females. Taxa that represent less than 0.5% of total sequences are clustered into “others”. In A., the multi-affiliated ASVs belonging to the *Pseudomonadota* correspond to multi-affiliated ASV of the *Moraxellaceae* and *Comamonadaceae* families and to the *Enterobacterales* order. In males **C**. and females **D**., genera above the threshold are represented in heatmaps showing their presence/absence in each individual ticks, and the mean infection rate in ticks are presented in %. **E**. Non-metric multidimensional scaling (NMDS) plot of the tick microbiota on the ASV level. The closer the points, the more similar microbial community composition. **F**. Heat map of the top 25 ASV that contribute the most to the difference between males and females.

At the genera level, nine and five genera above the threshold were identified in males and females. The five common genera were *Midichloria*, *Francisella*, *Rickettsia*, *Corynebacterium* and a multi-affiliated ASV (phylum *Bacillota*, class *Bacilli*). Almost all these genera were more encountered in males than females (**Figure 1A, 1B**), except for *Midichloria* whose number of sequences was 1.9 times higher in females (51.9%) compared to males (27.9%). *Francisella*, *Midichloria* and *Rickettsia* were present in 100% of both males and females, whereas *Corynebacterium* was present in all males but 95.3% of females, and the multi-affiliated ASV (class *Bacilli*) was present in all males and females except two of them (**Figure 1C, 1D**). There were four other genera above the threshold in the case of males, *Staphylococcus*, *Mycobacterium*, the multi-affiliated ASV of the *Bacilli* class and of the *Moraxellaceae* family and finally *Williamsia* (**Figure 1A**). They were not present in all males, but more encountered in males than females (**Figure 1C, 1D**).

The richness was significantly lower in females than males, as well as for the Shannon index, indicating that the alpha diversity was lower in females (**Figure S2, Table 1**). The NMDS analysis based on the Bray-Curtis dissimilarities showed the bacterial communities were clustered by tick sex (**Figure 1E**), as confirmed with the PERMANOVA test (p-value = 0.001, R^2^ = 0.22). The R^2^ value shows 22% of the variance in bacterial communities was explained by the tick sex variable. From the MOFA analysis, we generated a heatmap to visualise the 25 genera that make the biggest contribution to the difference between males and females. The heatmap was based on the weights from Factor 1, which captured 72% of the variance in the ASV dataset (**Figure 1F**). Most of the ASVs belonged to either *Francisella*, *Midichloria* or *Rickettsia*. The SIMPER analysis identified a total of 15 ASVs that significantly contributed to the Bray-Curtis dissimilarity metric between males and females (**Table S1**). Among them, 11/15 genera were affiliated to the *Pseudomonadota* phylum, they all together contributed to 59.2% of the difference (SIMPER contribution), but two genera in particular explained the most part of difference: *Midichloria* (35.0%) and *Francisella* (21.8%). There were 7/15 ASVs that were affiliated to the same genus; two were affiliated to *Rickettsia*, three to *Midichloria* and two to *Francisella*. Three genera of the *Bacillota* phylum also explained some of the difference, especially the multi-affiliated ASV of the *Bacilli* class (5.4%) (**Table 2**).

**Table 1:** Alpha diversity in males and females.

| Tick sex | N | Richness | p-value | Shannon | p-value | Inverse Simpson | p-value |
| --- | --- | --- | --- | --- | --- | --- | --- |
| Female | 128 | 39.4 ± 3.3% | 0.015 | 1.4 ± 3.6% | 0.006 | 2.7 ± 4.1% | 0.491 |
| Male | 153 | 45.7 ± 2.9% |  | 1.7 ± 4.1% |  | 3.4 ± 7.7% |  |

Mean observed values are shown for richness, Shannon and Inverse Simpson indexes. P-values (p) were calculated using the statistical models.

**Table 2:** Alpha diversity according to months for both males (n= 153) and females (n=128).

|  | Females |  |  |  |  |  | Males |  |  |  |  |  |
| --- | --- | --- | --- | --- | --- | --- | --- | --- | --- | --- | --- | --- |
| Month | Richness | p | Shannon | p | InvSimpson | p | Richness | p | Shannon | p | InvSimpson | p |
| February | 31,00 | 3.65<br>1x10<sup>-6</sup> | 1,23 | 0.43<br>86 | 2,47 | 0.63<br>31 | 45,71 | 5.16<br>6x10<sup>-8</sup> | 1,66 | 0.02<br>255 | 2,88 | 0.01<br>16 |
| March | 44,13 |  | 1,38 |  | 2,73 |  | 42,00 |  | 1,82 |  | 4,21 |  |
| April | 39,82 |  | 1,34 |  | 2,64 |  | 42,58 |  | 1,59 |  | 3,19 |  |
| May | 40,29 |  | 1,29 |  | 2,55 |  | 40,00 |  | 1,70 |  | 3,32 |  |
| June | 41,47 |  | 1,48 |  | 2,90 |  | 38,57 |  | 1,37 |  | 2,34 |  |
| July | 36,13 |  | 1,33 |  | 2,58 |  | 48,68 |  | 1,83 |  | 3,80 |  |
| August | 32,76 |  | 1,27 |  | 2,60 |  | 52,65 |  | 1,64 |  | 3,06 |  |
| September | 37,20 |  | 1,41 |  | 2,89 |  | 50,00 |  | 2,07 |  | 4,09 |  |

Mean observed values are shown for richness, Shannon and Inverse Simpson indexes. P-values (p) were calculated using the statistical models.

### Global temporal patterns in microbial networks of males and females

From the GLM model, we generated one male and one female network **(Figure 2A and 2B)** reflecting the monthly variations in microbial communities from February to September. The male and female networks differed in terms of the number of nodes, the composition of clusters, and their overall structure within the network. A total of 19 nodes were shared between the two networks. The female network was more diverse and spread out in terms of both node and edge numbers, while the male network was more compact, with a few dominant clusters (such as cluster 1M) playing a larger role in shaping the structure of the network.

**Figure 2:**
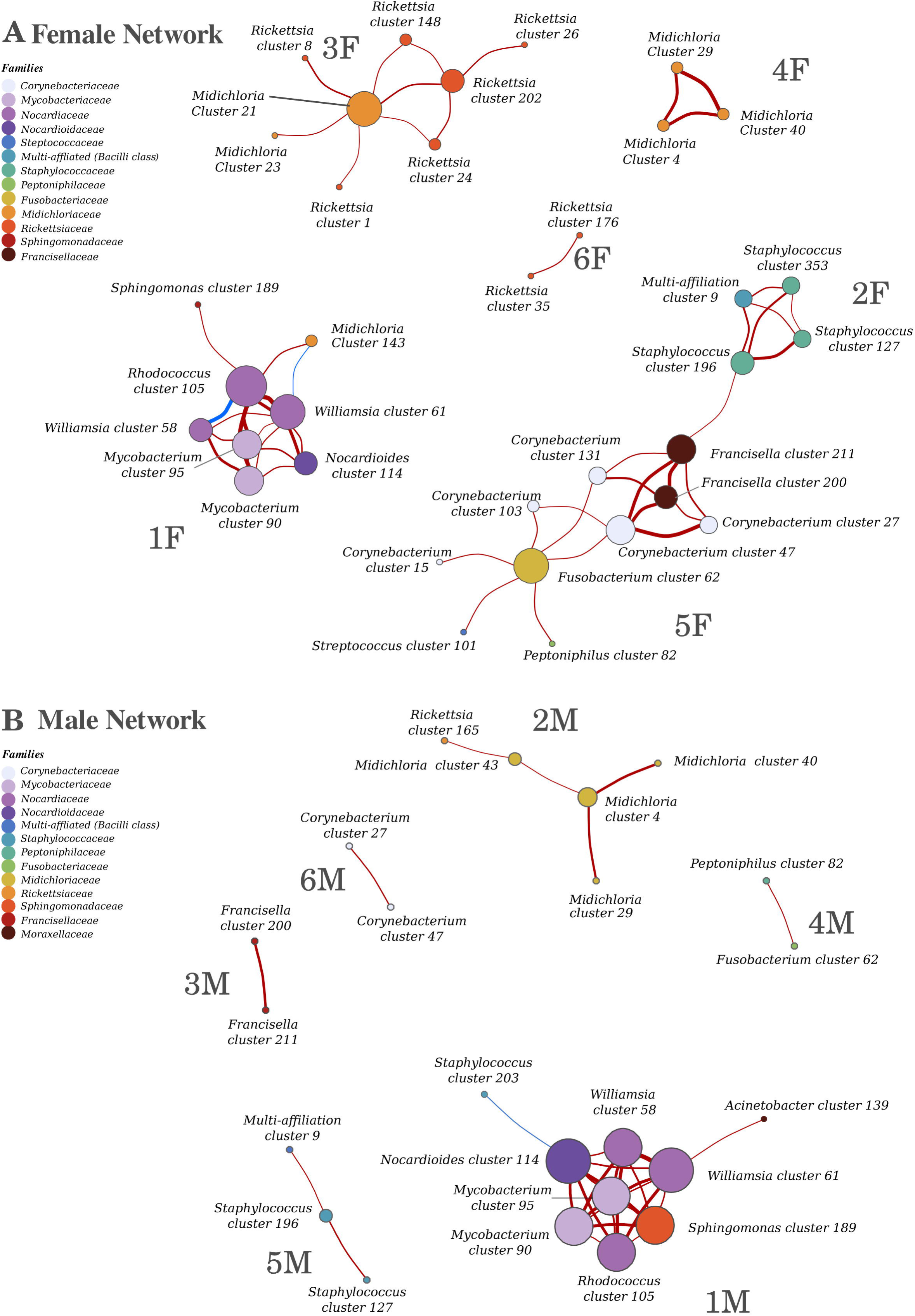
Static bacterial network based on **A**. the female dataset and **B**. the male dataset. Circles are coloured by taxonomic families. Each circle is a node or ASV. Here, a cluster refers to an ASV. Their size is proportional to the sum of the incoming edge weights. Thickness of the edge is proportional to the strength of the interaction. Positive interactions are represented by red edges, negative interactions are represented by blue edges.

The female microbial network (**Figure 2A**) had 35 nodes and 52 edges, it was composed of six unconnected networks (annotated as follow: 1F to 6F). Edges with a high weight represented a dominant influence or an important ecological relationship, such as a key mutualistic or antagonistic interaction that significantly affected the microbial community structure or function. Nodes with higher cumulative interaction strengths (sum of the absolute weights of all connected edges) were larger, indicating that they are involved in stronger or more significant correlation within the network. Out of the 52 edges, two relationships had negative weights: one between *Rhodococcus* ASV_105 and *Midichloria* ASV_143 in cluster 1F (weight = -0.027239434), and another between *Williamsia* ASV_61 and *Midichloria* ASV_143 in cluster 1F (weight = -0.004617648). These negative interactions were highlighted in blue in **Figure 2A**. It means that an increase in one species was associated with a decrease in the other. The cluster 1F comprised eight environmental ASVs belonging to *Mycobacterium* (2 ASVs), *Williamsia* (2), *Rhodococcus*, *Nocardioides, Sphingomonas* genera, and one *Midichloria* ASV. It was a densely connected cluster, almost fully connected with bacteria having strong interaction with each other. Cluster 2F was composed of four ASVs including three *Staphylococcus* ASVs and a multi-affiliated ASV of the *Moraxellaceae* family. Cluster 2F consisted of four ASVs, including three *Staphylococcus* ASVs and one multi-affiliated ASV from the *Moraxellaceae* family. The cluster 3F was composed of six *Rickettsia* ASVs and two *Midichloria* ASVs, while cluster 4F comprised only three *Midichloria* ASVs. The large node size of *Midichloria* ASV_21 suggested its interactions had strong weights across its five connections to *Rickettsia* (ASVs 1, 8, 21, 24 and 202), indicating that *Midichloria* played a central or dominant role within this sub-network. This species had strong and positive interactions affecting the surrounding bacteria. The cluster 5F comprised eight environmental ASVs including five *Corynebacterium* ASVs, one *Streptococcus*, one *Peptoniphilus* and one *Fusobacterium* ASV, as well as two *Francisella* ASVs. This cluster was linked to cluster 2F by a weak interaction between the ASV *Francisella*_211 (cluster 5F) and the ASV *Staphylococcus*_196 (cluster 2F). Based on the size of the node, *Fusobacterium* ASV_62 strongly affected: *Peptoniphilus, Streptococcus and Corynebacterium*, and based on the edges weights, we noticed a high strong ecological relationship between *Francisella* and *Corynebacterium*. Finally, the cluster 6F was composed of two *Rickettsia* ASVs.

The male microbial network has 23 nodes and 32 edges, it was composed of six unconnected networks (annotated as 1M to 6M). Out of the 32 edges, one relationship has negative weights: *Nocardioides* ASV_114 and *Staphylococcus* ASV_203 in the cluster 1M (weight -0.019438662) (**Figure 2B**). The cluster 1M was composed of the same seven ASVs observed in female cluster 1F, with the addition of two ASVs (*Acinetobacter*_139 and *Staphylococcus*_203). Based on the connectivity, edges and nodes weights, we observed that the bacteria in the cluster 1M have a strong mutualistic relationship and compared to the other clusters in the male network, this cluster played a key role in the male microbiome. The cluster 2M included the same ASVs of the cluster 4F, except that it comprised two additional ASVs (*Midichloria*_43 and *Rickettsia*_165), which were not in the cluster 4F. The remaining male clusters (3M to 6M) consisted of few nodes, with each cluster containing only two to three nodes. The cluster 3M was composed of two *Francisella* ASVs, while the others were composed of environmental ASVs including *Peptoniphilus* and *Fusobacterium* (cluster 4M), two *Staphylococcus* and a multi-affiliated ASV of the *Moraxellaceae* family (cluster 5M) and finally two *Corynebacterium* ASV (cluster 6M).

### Temporal shifts in male and female microbiota across the months Females

The temporal patterns were analysed separately for males and females. The richness significantly varied with months (p-value = 3.651×10^−6^) while the Shannon and Inverse Simpson indexes did not (**Table 2**, **Figures S1**). The richness was higher in winter compared to summer (p-value = 0.0002). The PERMANOVA test showed a significant contribution of the month to the Bray-Curtis dissimilarities, (p-value = 0.018), the R^2^ value indicated that 9.4% of the variance in bacterial communities was explained by the month. Additionally, the NMDS analysis supported this effect, as the ordination plot displayed a distinct influence of the month on the arrangement of bacterial communities (**Figure 3A**). The stress value was lower than 0.2, which indicated a limited risk of pattern misinterpretation and confirmed the robustness of the NMDS analysis.

**Figure 3:**
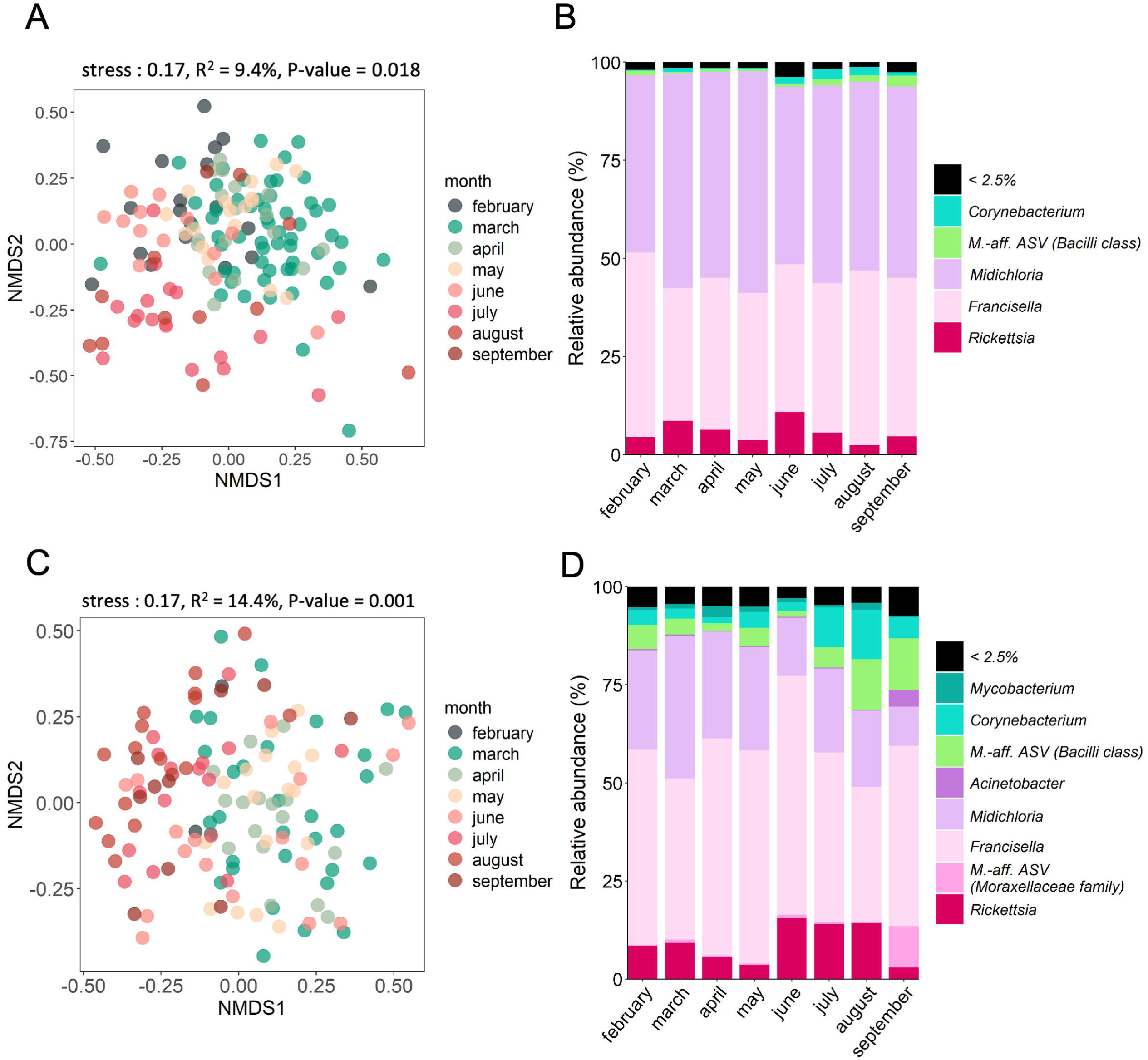
Temporal dynamics of *H. marginatum* microbiota for female (upper panel) and males ticks (bottom panel). **A**. and **C**. NMDS plots of bacterial communities based on Bray-Curtis dissimilarity among tick samples. **B**. and **D**. Barplots summarising the relative abundance (%) of the most abundant genera (>2.5% of monthly relative abundance).

The influence of months on the relative abundance of the most abundant genera (those presented in **Figure 3B**) showed that the month did not significantly influence *Francisella*, *Midichloria*, *Rickettsia* and *Acinetobacter* but did influence *Corynebacterium*, the multi-affiliated ASV class *Bacilli*, the multi-affiliated ASV of the *Moraxellaceae* family and *Mycobacterium* (**Figure 3B, Figure S3A, Table S2**).

We noticed, 16 genera were absent for at least one month, including *Trueperella*, *Streptococcus*, *Sphingomonas*, *Rhodococcus*, *Pseudomonas*, *Porphyromonas*, *Peptoniphilus*, *Nocardioides*, *Mannheimia*, *Helcococcus*, *Halomonas*, *Comamonas*, *Chryseobacterium*, *Caviibacter*, *Caenimonas*, *Brevundimonas* (**Figure S4A**). Of these, 11 were absent for one month, four were absent for two months and one for three months. A higher number of absent genera was noticed in February (10 genera) compared to April (4 genera), August (3 genera), September (3 genera) and July (2 genera). All these genera did not represent a major part of the microbiota since they represented between 0.01% and 0.07% of the total number of sequences in females.

As the genera comprised several ASVs, the variation in their relative abundance was represented in **Figure S5** and **Figure S6**. The range of relative abundance variation across the eight months for all ASVs belonging to the *Actinomycetota* phylum was 7.2%, 10.8% for *Pseudomonadota*, 14.5% for *Bacillota*, 15.3% for *Bacteroidota* and 18.2% for *Fusobacteriota*. Strong monthly variations were observed for the relative abundance of some specific ASVs such as *Corynebacterium* ASV 15, 47, 103, whereas the other *Corynebacterium* ASVs (129, 131, 25, 27, 88) did not exhibit such variations (**Figure S5**).

### Males

The alpha diversity significantly varied with months for the richness (p-value = 5.166×10^−8^), the Shannon index (p-value = 0.02255) and the Inverse Simpson index (p-value = 0.0116) (**Table 2**, **Figures S2**). For example, the richness was higher in spring (p-value = 0.0953) and winter (p-value = 0.0001) compared to summer months. The PERMANOVA test showed significant contribution of the month to the Bray-Curtis dissimilarities (p-value = 0.001), the R^2^ value indicated that 14.4% of the variance in bacterial communities was explained by the month. The NMDS analysis based on the Bray-Curtis dissimilarities showed that the bacterial communities were slightly clustered by month (deduced by graphic reading) (**Figure 3C**). As observed for females, the stress value that was lower than 0.2 represents a good ordination with little risk of pattern misinterpretation.

We analysed the influence of the month on the relative abundance of the most abundant genera (those presented in **Figure 3D**). On those eight genera (*Francisella*, *Midichloria*, *Corynebacterium, Mycobacterium*, the multi-affiliated ASV class *Bacilli*, the multi-affiliated ASV family *Moraxellaceae* and *Acinetobacter*), the month did influence their relative abundance, except for *Rickettsia* for which no influence was observed (**Figure 3D, Figure S3B, Table S2**).

We noticed 11 genera were absent for at least one month, including *Williamsia*, *Trueperella*, *Pseudomonas, Porphyromonas, Peptoniphilus, Nocardioides*, a multi-affiliated ASV (class *Bacilli*)*, Mannheimia, Helcococcus, Caviibacter* and *Caenimonas*. Nine were shared with females including *Trueperella, Pseudomonas, Porphyromonas, Peptoniphilus, Nocardioides, Mannheimia, Helcococcus, Caviibacter* and *Caenimonas*. Of the 11 absent genera in males, six were absent for one month, four were absent for two months and one for three months (**Figure S4B**). A higher number of absent genera was found in September (10 genera), followed by April (3 genera), June (2 genera), July (2 genera) and February (1 genera). Some of these genera constituted a minor proportion of the microbiota, including *Porphyromonas*, *Helcoccoccus*, *Caviibacter*, *Caenimonas* and *Mannheimia* (between 0.01% and 0.07% of the total sequences in males). Conversely, other genera represented a more substantial proportion, including *Williamsia, Nocardioides, Trueperella, Peptoniphilus, Pseudomonas* and a multi-affiliated ASV (family *Comamonodaceae*) (between 0.17% and 0.94% of total sequences in males).

As genera comprised several ASVs, the variation in their relative abundance was represented in **Figure S5** and **Figure S6**. The range of relative abundance variation across the eight months for all ASVs belonging to the *Actinomycetota* phylum was 34.6%, 31.7% for *Fusobacteriota*, 23.6% for *Bacillota,* 17.3% for *Bacteroidota* and 17.2% for *Pseudomonadota*. Strong monthly variations were observed for specific ASVs, such as *Corynebacterium* ASVs_25, 15 and 103 (**Figure S5**).

### Networks

The analysis of the female and male networks across the months revealed significant changes into the composition and number of nodes, edges, and predominant nodes (larger node) (**Figure 4**, **Figure 5**). For instance, in females, in May, the larger node was *Rhodococcu*s_105 whereas in June, *Rhodococcus*_105 was a small node and the larger one was *Fusobacterium*_62, which disappeared from the network of July, August and September, for which larger nodes were respectively *Corynebacterium*_103, *Corynebacterium*_131 and *Staphylococcus*_353 (**Figure 4**). The predominant node also changed along with the month in males (**Figure 5**). For instance, it was *Midichloria*_161 and *Pseudomonas*_94 in February while the next month, *Midichloria*_161 was a very little node. The following months, in April, the biggest nodes were *Corynebacterium*_88 and once again a *Pseudomonas* ASV but a different one compared to February (ASV 54) while in May, it was *Midichloria*_4 and *Staphylococcus*_203 that were predominant. In June, the biggest clusters were *Francisella*_5 and the multi-affiliated ASV of the *Moraxellaceae* family, *Fusobacterium*_62 in July, and *Williamsia*_61/ *Rhododoccus*_105 in August. In summer, especially for June and July, the networks were less complex (number of interactions) and less structured compared to winter and spring months (from February to May). By August, the network became more robust, with several ASVs, including *Midichloria*_43, *Rhodococcus*_105, *Williamsia*_61, and *Xanthomonas*_83.

**Figure 4:**
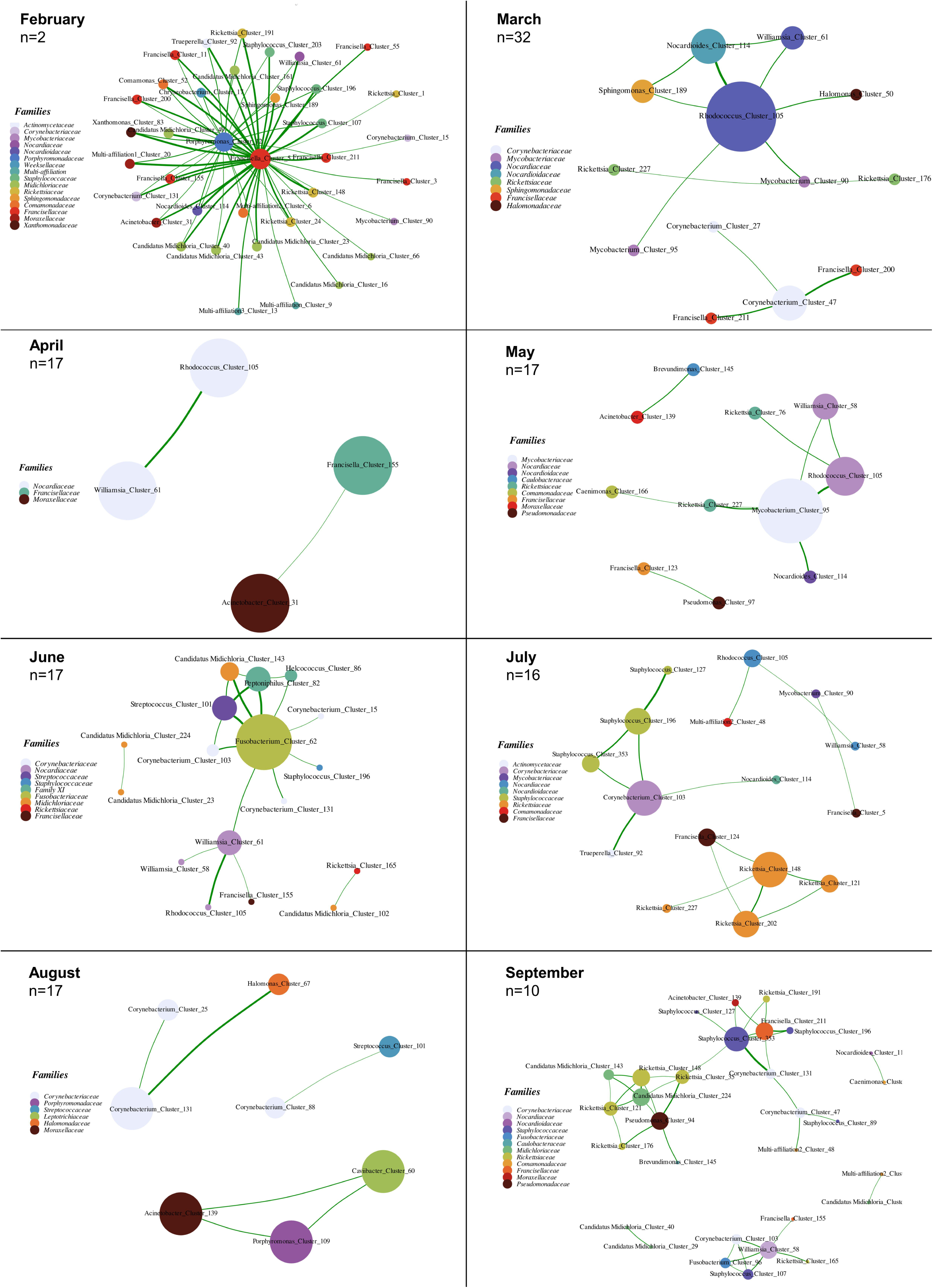
Monthly subnetworks of female ticks over the year from February and September. Number of ticks is indicated in the upper left corner of each panel that delimitates months. The colours of nodes are indicative of the family taxonomic rank of ASVs, they are inherent to each panel.

**Figure 5:**
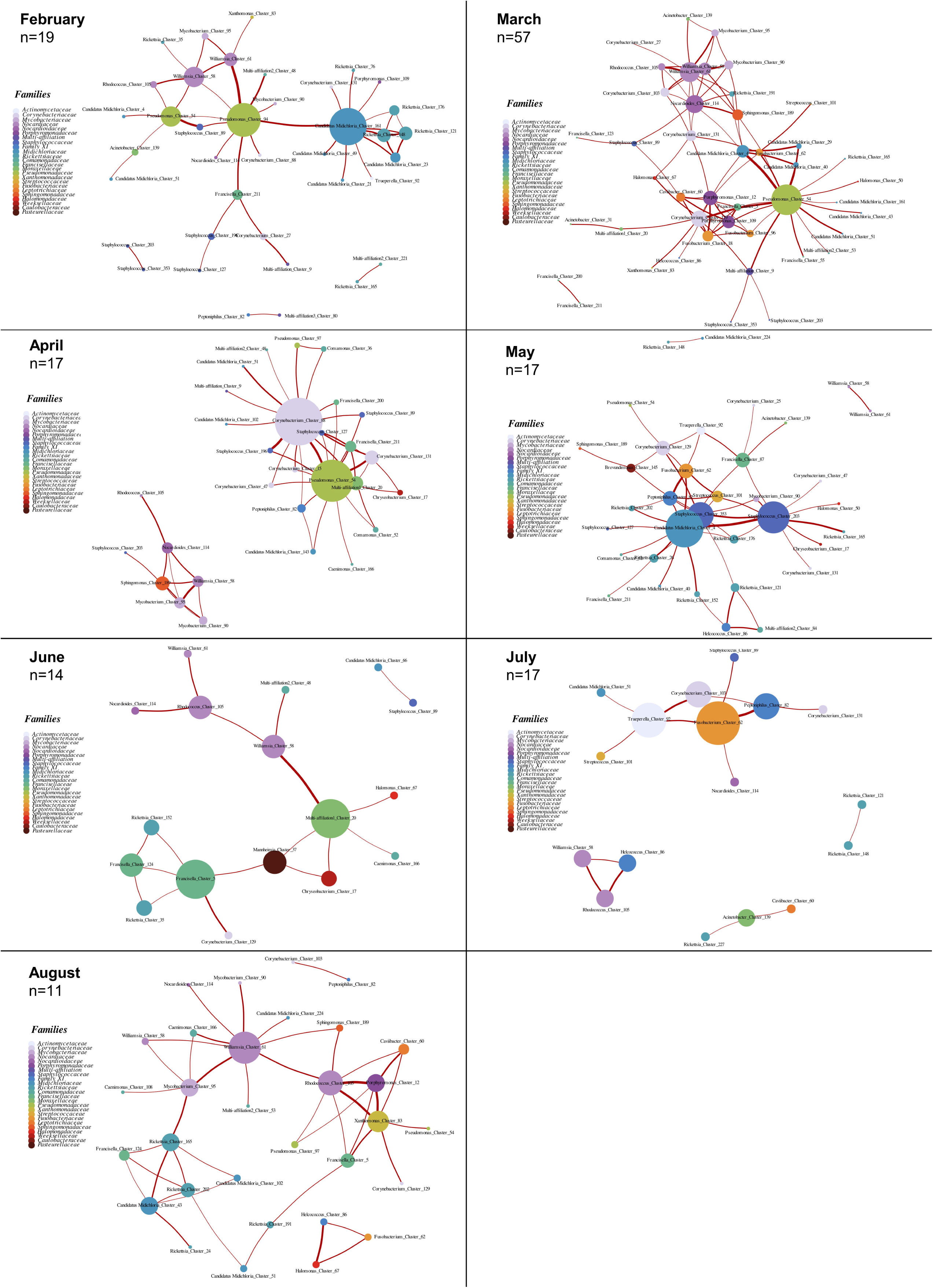
Monthly subnetworks of male ticks over the year from February and August. Number of ticks is indicated in the upper left corner of each panel that delimitates months. The colours of nodes are indicative of the family taxonomic rank of ASVs, they are inherent to each panel. September network is not shown because of not enough individuals (n=1).

If we specifically focus on the interactions between *Midichloria* and *Rickettsia* throughout the month for tick females, we notice that direct interactions between two ASVs of these genera only appeared in the networks of June and September. Interestingly, there were no other interactions, even indirect ones, during the other months. In males, both direct and indirect interactions were observed more regularly throughout the sampling period. Indeed, direct interactions appeared in February, May, and August. During the months that lacked direct interactions, indirect interactions between *Midichloria* and *Rickettsia* ASVs were detected in March with *Pseudomonas* as an intermediate node. This focus on the *Rickettsia* - *Midichloria* interactions showed that these interactions are not constant across the months particularly in females.

When comparing the evolution of edges/interactions through time between both male and female microbial communities, the male networks presented a higher number of interactions compared to the female ones throughout the months (**Figure 4, 5**). In addition, these interactions formed a more complex structure, a tighter network than for females. In other words, the male networks were more structured than the female ones throughout the months. The nodes (ASVs) involved in the networks of males were more stable over time. This information is synthesised in the **Figure S7**, which shows the female networks presented more frequent reorganisation and movement among nodes across time, a highly variable number of interactions and several clusters that appeared and changed overtime. Conversely, the male networks exhibited a more organised structure, with more stable node composition, a more stable number of interactions over time, therefore the clusters were more stable overtime, more consistent. For instance, some nodes, such as *Pseudomonas*_54, remained consistent with high cumulative weights, particularly in February, March, and April.

## Discussion

### Global description of the *H. marginatum* microbiota

The objective of this study was to identify the factors that shape the holobiont *Hyalomma marginatum*, its microbiota and its interactions, with a particular focus on the influence of the tick sex and the temporal dynamics. *H. marginatum* is a tick species of interest in public health, given its role as a vector and reservoir for CCHFV, the only pathogen formally identified to be vectorized by this tick species (Bernard *et al*. 2024b; Bonnet *et al*. 2023). It has been demonstrated it can carry other pathogens belonging to the *Anaplasma*, *Theileria* and *Rickettsia* genera. However, its role as a vector for these pathogens remains unknown (Bernard *et al*. 2024a; Joly-Kukla *et al*. 2024a, b). An understanding of the temporal dynamics of the *H. marginatum* holobiont will enhance the general knowledge base concerning the factors that influence the microbiota, which may have a substantial impact on the tick vector competence.

We showed that the *H. marginatum* microbiota was dominated by *Francisella* and *Midichloria*, together representing a major part of the total number of sequences in both males (74.7%) and females (89.6%). These bacteria were present in 100% of individual ticks analysed. This result was obviously expected because these bacteria are known to form a dual-partner nutritional system (co-symbiosis) that provide B vitamins essential for the survival of *H. marginatum* (Azagi *et al*. 2017; Buysse *et al*. 2021). A third bacterial genera, *Rickettsia sp.*, was extensively detected in ticks. Our sequencing approach did not allow us to directly obtain the affiliation of taxon at the species level. However, we can clearly state that most of sequences correspond to the species *Rickettsia aeschlimannii*, specifically identified in the same samples using two other different detection techniques including a specific qPCR and a real-time PCR (high-throughput screening using microfluidics) (Joly-Kukla *et al*. 2024a). Based on different arguments including the maternal transmission, the high infection rates and its strong spatial and temporal variations, we previously suggested that *R. aeschlimannii* might be a secondary symbiont in *H. marginatum* (Joly-Kukla *et al*., 2024a, b). All ticks were positive for *Rickettsia* which is highly consistent with the results of Azagi *et al*., (2017) that demonstrated 95% of *H. marginatum* ticks collected from Israel were positive for *R. aeschlimannii*. In contrast, our previous studies demonstrated that this bacterium was detected in only 45% of the same tick samples analysed here. These contrasting results observed on the same ticks could be explained by a first hypothesis involving a technical limitation, for example, a certain bacterial density threshold that would be required to successfully detect the bacterium when using the microfluidics method. This hypothesis would be in line with the high maternal transmission rate that is observed for *R. aeschlimannii* (Azagi *et al*., 2017). A second significant hypothesis implies that some sequences belonging to the genus *Rickettsia* detected in the present study might correspond to different *Rickettsia* species like *R. raoultii*, a very close species of *R. aeschlimannii*.

Many microbiota members identified in this study, including *Francisella*, *Midichloria*, *Rickettsia*, *Corynebacterium* and *Staphylococcus*, were previously reported in another study that analysed the microbiota of 30 *H. marginatum* ticks collected from cattle between June and August 2020 in different geographic location of Tunisia, using the same sequencing methodology (Illumina Miseq) but not exactly the same target region (v3-v4 region 16S rRNA gene) (Benyedem *et al*. 2022). These genera were also detected in a second study utilising the same sequencing methodology (Illumina, V4 region 16S rRNA gene) on 23 *H. marginatum* ticks collected from cattle in Corsica (Maitre *et al*. 2023). This result suggests that bacterial members of the *H. marginatum* microbiota are shared independently of spatial or temporal scales. In contrast, we did not identify certain bacterial members that were previously detected in the study of Maitre *et al*., (2023) including *Cutibacterium* and *Jeotgalicoccus*, which were previously identified as keystone bacteria (highly connected taxa driving community composition) in their study. More precisely, *Cutibacterium* sequences were considered as kit contaminants regarding our negative controls, or removed by the abundance ASV filters we applied. These results underline one more time the need for numerous negative controls when studying the tick microbial communities (Lejal *et al*. 2020) and might also be explained by the fact that Maitre and al., (2023) considered the tick’s cuticle bacterial communities as part of the tick’s microbiota and therefore did not perform a tick washing step before nucleic acid extraction, while we did.

Interestingly, among the most abundant bacterial genera in *H. marginatum* (**Figure 1A, 1B**), some are known to comprise species commonly associated with the environment, including *Staphylococcus*, *Corynebacterium*, *Williamsia*, *Acinetobacter* and *Mycobacterium* (Khoo *et al*. 2016; Portillo *et al*. 2019), and were previously detected in several tick species (*Rhipicephalus, Haemaphysalis*, *Dermacentor* or *Ixodes*). These bacteria were present in a significant proportion of individual ticks whether males or females (**Figure 1C, 1D**), and were found in interactions with each other in the two bacterial networks (**Figure 2**). In some clusters identified in both networks (*i.e.* clusters 1F and 1M in **Figure 2**), some of these bacterial genera presented strong mutualist relationships suggesting that they could play a significant role in the tick microbiome. *Staphylococcus* was found in 90.2% of males and 85,2% of females and represented 0.6% and 0,3% of sequences. This genus is frequently reported in several tick species of the following genera: *Ixodes*, *Hyalomma*, *Haemaphysalis* and *Rhipicephalus* (Abraham *et al*. 2017; Alreshidi *et al*. 2020; Hernandez, Salamat, et Galay 2023; Lim *et al*. 2020). It is even suggested to be a key member of the microbiota of *Rhipicephalus microplus* (Adegoke *et al*. 2020; Hernandez, Salamat, et Galay 2023). *Corynebacterium* was found in 100% of males and represented 4.4% of sequences. it was present in 95.3% of females and represented 1.2% of sequences. It was also reported in *I. scapularis* gut tissues (Abraham *et al*. 2017), and in other tick species as *Haemaphysalis*, *Dermacentor*, *Amblyomma* and *Rhipicephalus* (Lim *et al*., 2020, Hernandez *et al*., 2023). In *Amblyomma testudinarium* (n=13), the observation of high relative abundances for both *Corynebacterium* (8.8%) and *Staphylococcus* (21.7%) led the authors to suggest that these bacteria could have a role in the tick biology or the transmission of pathogens (Lim *et al*., 2020). In the present study, we suggest that *Staphylococcus* and *Corynebacterium* (**Figure 1**) might originate from either the horses’ skin, since they are common skin-associated bacteria (Baquer *et al*. 2023; Greay *et al*. 2018; Thanchomnang *et al*. 2023), or directly from the environment (Ray *et al*. 2022; Singh et Kumari 2023). These bacteria were likely ingested by ticks, since ticks were previously washed with bleach which normally aims to eliminate the cuticular bacteria (Binetruy *et al*. 2019b). Our results showed a high infection rate, a high relative abundance and strong interactions between these bacteria. Based on these results, we suggest that *Staphylococcus* and *Corynebacterium* might play a role in the colonisation of other microbes including pathogens. Additionally, it is interesting to note that four biotin genes were observed in low abundance in contigs of *Staphylococcus* in *H. marginatum* (Buysse *et al*. 2021), which lead to open questions: could these bacteria be involved in physiological processes in ticksIf so, is there a maternal transmission or a mixed transmission? The same questions arise for the three other abundant environmental-associated bacteria including *Williamsia*, *Mycobacterium* and *Acinetobacter*.

In both males and females, we noticed the clusters composed of environmental bacteria (**Figure 2**, Cluster 1F, 1M, 4M, 5M, 6M) were most of the time isolated from the cluster composed of endosymbiotic bacteria (**Figure 2**, 3F, 4F, 6F, 3M). This observation is consistent with the fact that bacteria are specialised and located in different organs. For instance, one of the most abundant bacteria was the obligate symbiont *Midichloria* known to be located preferentially in malpighian tubules and ovaries (Olivieri et al., 2019). Except for one ASV (*Midichloria*_143) that interacted with environmental bacteria (**Figure 2**, Cluster 1F), the five other *Midichloria* ASVs formed isolated clusters and interacted only with other *Midichloria* or *Rickettsia* ASVs (**Figure 2**, Cluster 3F, 4F). Since the environmental bacteria are likely to be mainly encountered in the tick midgut (Ross *et al*. 2018), a few of them naturally interact with endosymbionts, due to the limited overlap in their organ locations. The microbiota composition, including pathogens, typically vary according to the tick’s organs (Pollet *et al*. 2020), and encourages to include this scale into tick holobiont studies, but the experimental conditions for dissections can be challenging, and cross-contaminations are to be avoided. However, we observed in the female network that the obligate endosymbiont *Francisella*-LE interacted with environmental-associated bacteria including *Corynebacterium* and *Staphylococcus* (**Figure 2**, Clusters 5F, 3M, 6M). The large node sizes further imply a significant relationship between *Francisella*-LE and *Corynebacterium* in the female microbial communities. This interaction can suggest that some *Francisella*-LE bacteria were possibly located in the midgut, or that *Corynebacterium* and *Staphylococcus* were located in Malphighian tubules or ovaries, which would raise the question of their function in physiological processes and maternal transmission. Otherwise, in bacterial networks at the whole tick level, a co-occurrence (connection) between two ASVs can mean there is a direct interaction in the tick within the same tick organ, but it can be an indirect interaction from two different body locations. For instance, indirect interaction can occur through a bacterium that activates the immune system in a given organ and have repercussions in bacteria of other tick locations (*i.e.* Aivelo *et al*., 2019). In the bacterial network of males and females (Figure 2), we reported positive interaction between *Rickettsia* and *Midichloria* ASVs. Interactions between these two genera were reported in the literature for *I. ricinus* (Lejal *et al*. 2021), as well as *A. maculatum* between *R. parkeri* and *Midichloria mitochondrii* (Adegoke *et al*. 2022; Budachetri *et al*. 2018b). This interaction could be direct by involving a specific metabolite that is consumed by the other, or indirect by an action on the immune system by one partner that favours the other (Gall *et al*. 2016). Further investigations will be needed to identify the molecular drivers of the interaction, the potential effects in the tick physiology, by analysing the transcriptome of ticks infected or not by these two bacteria, and the transcriptome of the bacteria themselves.

### Variation in microbiota composition and interactions according to the tick sex

The *H. marginatum* bacterial communities varied according to the tick sex. The female microbiota was less diverse, as already demonstrated in several tick species including for example *Dermacentor variabilis* (Duncan *et al*. 2022), *Amblyomma Americanum* (Trout Fryxell et DeBruyn 2016), *Ixodes scapularis* and *I. affinis* (Krawczyk *et al*. 2022; Thapa, Zhang, et Allen 2019; Van Treuren *et al*. 2015). In their study, Gofton *et al*., (2015) demonstrated that a technical bias can occur during the sequencing process when abundant endosymbiotic bacteria such as *Midichloria*, which dominates the microbiota of *Ixodes* ticks and recruits more reads. This leads to the masking of less abundant bacteria and pathogens that are less detected. In our study, we observed that the number of sequences belonging to the two primary endosymbionts *Francisella*-LE and *Midichloria* was lower in males (167,000 sequences) than females (199,125), demonstrating a dominance of endosymbionts in females. This is in line with the statistical analysis that showed these two obligate endosymbiont genera together contributed to 56.8% of the dissimilarity between males and females (Bray-Curtis dissimilarity) (SIMPER analysis, **Table S1, Figure 1F**). Based on these results and in accordance with the hypothesis proposed by Krawczyk *et al*., (2022), we postulate that endosymbiotic bacteria are more abundant in females, concealing the less abundant ones in comparison to males. This provides a potential explanation for the observed difference in alpha diversity. The biological interpretation of endosymbionts being more abundant in females can be linked to the lack of ovaries in males, an organ where the maternal transmission occur in germinal cells and thus associated with higher bacterial multiplication (Christensen *et al*. 2019; Daveu *et al*. 2021; Duron *et al*. 2018; Olivieri *et al*. 2019; Vautrin et Vavre 2009). Endosymbionts observed in males are likely located in Maphigian tubules whereas in females, they are spread between Malpighian tubules and ovaries. In addition, males usually do not feed as much as females, which makes nutritional endosymbionts less necessary (Bonnet *et al*. 2017). This biological interpretation is supported by the comparison between male and female networks, where the sizes of *Midichloria* and *Francisella* nodes were smaller in the male network compared to the female one (**Figure 2**). This means that these nodes presented a lower influence in males suggesting that these endosymbionts would probably be less involved in the structure of the male microbiota network.

### Temporal dynamics in *H. marginatum* microbial communities

Since the tick sex was an important factor shaping the *H. marginatum* microbiota, we analysed the temporal dynamics separately for males and females. Temporal patterns were analysed at the level of the month, from February to September 2022. When looking at the scale of entire bacterial communities, the month and the season did have a significant influence on alpha diversity, while the richness, Shannon and Inverse Simpson index were significantly influenced by the month for males, only the richness was influenced in the case of females. The richness was higher in winter compared to summer for both males and females, which means more species were observed in ticks collected in winter compared to those collected in summer. The beta diversity was also influenced by the month for both males and females. In the PERMANOVA analysis of Bray-Curtis dissimilarities, the month explained 9.4% of the model variance for females and 14.4% for males. Based on the NMDS **(Figure 3A, 3C**), we distinguished two major groups: ticks collected in March/April, and those collected in July/August. These results suggest that biotic or abiotic factors may drive the microbiota composition through the time. For example, during its non-parasitic stages in the environment (floor litter), the diversity of environmentally-acquired bacteria could be influenced by the different microbiota of floor litter that changes throughout the months and seasons due to climate and temperature. Additionally, the temperature could affect the tick’s behaviour, which can hide into the floor litter to escape from too warm temperatures (Stachurksi *et al*. comm. pers.), thus confronting the tick to different bacteria than at the surface of the floor. During its parasitic stages on hosts, the tick microbiota diversity could also be affected by the microbiota of the host skin, which can change because of sweat for example. The temperature has already been identified as a key factor influencing the microbiome of *Ixodes ricinus*, as it decreased the alpha diversity and affected the relative abundances of several bacterial genera (Thapa *et al*. 2019). These hypotheses on the acquisition of different environmental bacteria depending on the time in the year are consistent with our results reporting that the relative abundance of environmental genera was statistically influenced by the month (**Table S2**). About 20% of the total genera of females were absent from all ticks collected in at least one month, which underlines there is a significant proportion of genera occurrence that is variable among months (**Figure S4**). For example, *Porphyromonas* was absent from all the females of April and July and from all the males in June and July. At an even finer scale, it was interesting to note that for some genera comprising several ASVs, only a fraction of these ASVs was actually variable over months. For instance, 5/8 *Corynebacterium* ASV relative abundances were actually variable over months. This result allows us to hypothesise that the temporal variations occur at a finer scale than the genus, at the species level and maybe the strain level. These ASV-specific dynamics underline the limitations of the genus resolution using 16S rRNA sequencing.

In females, among the top 8 most abundant bacterial genera (**Figure S3, Table S2**), only the relative abundance of 4/8 genera did vary over time, they were all environmental bacteria. The endosymbionts were stable in females, whereas for the males, 7/8 of the most abundant bacteria were significantly influenced by the month, including both the endosymbionts and environmental bacteria. This result suggests that the temporal stability of the female microbiota is higher. This might be explained by the need of endosymbionts for females since they feed throughout the whole activity period between February to September, independently from the month, while males feed little or not at all, and therefore could not need as much as females on nutritional endosymbionts.

For *Rickettsia*, even though at the scale of the genus there were not any differences, strong variations over months were nevertheless observed at the scale of the ASVs. In a previous study, we identified strong variations of its infection rates in the same collected ticks, using *OmpB* gene and *glta* gene. Here, we noticed 100% of these ticks were positive for *Rickettsia*. As already explained, the fact we did not detect *R. aeschlimannii* in 100% of ticks in our previous study while this is the case here, might be explained by a certain bacteria density threshold that prevents the detection using the microfluidics method. The temporal variation of infection rates using microfluidics might be a reflection of bacteria abundance / density. To support this hypothesis, in this paper, while no statistical differences were observed for its relative abundances, we noticed the same temporal pattern from February to September that we previously reported using the Biomark assay. Even if bacteria are present in 100% ticks independently from the month, the bacterial loads might be higher in specific months compared to the others, and coupled with the threshold hypothesis, it might be reflected by the infection rates.

### Interaction patterns vary across the months

As observed more generally by Deutschmann *et al*., (2023), several studies dealing with tick microbial community interactions have used temporal microbial abundance data to infer static networks summarising all potential interactions in space and time (Aivelo, Norberg, et Tschirren 2019; Lejal *et al*. 2021; Maitre *et al*. 2023; Mateos-Hernández *et al*. 2021; 2020). This static abstraction assumes that the network topology does not change and edges represent persistent interactions assumed as interactions; that is, edges are present throughout time and space. This assumption cannot represent the reality of most microbial interactions. This means that we need to investigate the dynamics of microbial interactions in ticks. This is particularly pertinent given that recent studies suggest the use of anti-microbiota vaccines based on the detection of keystone bacteria determined by static networks (Aželytė *et al*. 2022; Mateos-Hernández *et al*. 2021; 2020). In our study, we demonstrated that the bacterial networks of both males and females were highly dynamic and variable over the months. All the network parameters, the composition and the number of nodes, the network centrality, the number and the type of edges and the global structure varied for both males and females. Both **Figures 2** and **4A** demonstrate how the notion of scale is crucial to consider when studying potential interactions in tick microbial communities using network analysis. For example, we observed in **Figure 4A** that the interactions between *Rickettsia* and *Midichloria* ASVs were not always reported across the months and were only present in June and September for females, and in February, May and August for males. In contrast, we noticed that this cluster of interactions was an important part of the female network throughout all the months (**Figure 2**). This example illustrates the influence of the scale at which we study tick bacterial interactions and how it can change the results.

In a complementary way, our findings also demonstrate that contrasting temporal patterns in tick microbial interactions were observed between males and females. Overall, by comparing monthly networks, we observed that the male networks presented a higher number of interactions than for females, they were more structured, this structure being more stable over time. The interactions were more consistent and the node composition more stable over time compared to the female networks. For females, the networks were less structured, they showed frequent changes into the node composition and variations in the number of interactions over time. The temporal evolution of the female networks indicates a more complex and rapidly changing environment, whereas the male networks were more consistent. A solid biological interpretation of our results is difficult at this stage but this information is crucial in regards to new tick control strategies recently proposed and targeting key members of the tick microbiota. As an example, anti-microbiota vaccines have recently been suggested and tested to remove ubiquitous and abundant keystone bacteria and finally alter the tick physiology and vector competence (*i.e.* Mateos-Hernández *et al*., 2020, 2021). Such an approach necessarily implies that both the presence and interactions of these keystone bacteria with the other members of the tick microbiota are stable over time. In regard to our results, we thus call for caution when keystone bacteria are identified in tick microbial networks since we demonstrated that bacterial interactions greatly vary according to the scale of the study, at the level of the tick sex and at the monthly scale. As an example, *Staphylococcus* was identified as a keystone taxon in *Rhipicephalus bursa* networks (Maitre *et al*., 2023), and proposed as a candidate for anti-microbiota vaccine to decrease the *Rickettsia* transmission in *R. bursa*. The use of microbiota for vaccines is a promising approach for innovant control strategies that still need investigations which in this example, would require a temporal study of *R. bursa* networks, in order to characterise if *Staphylococcus* represent a keystone bacterium at any time of the year. If not, the efficacy of a vaccine would be highly limited to specific periods in the year, when this keystone taxon is present and central in the networks.

Network analysis is a powerful tool to unravel microbial interaction within ticks, but more investigations are needed at different scales. In our case, it would be necessary to know if monthly patterns are stable over years. In addition, our study focused on the whole tick scale and even if our network analyses generally distinguished interactions between bacteria known to be localised either in ovaries-malpighian tubules or in the midgut, networks should also be investigated at the scale of the tick organs in order to gain insights into the direct microbe-microbe interactions. Spatial investigations on the effect of the site, the region, the population genetics on microbial networks are also needed to collect information about keystone bacteria and the specific context in which anti-microbiota vaccines could be applied and actually work.

## Declarations

## Acknowledgment

The authors are grateful to Frédéric Stachurski from UMR ASTRE or his contribution in the fieldwork. Thank you to everyone involved in the HolisTique project for ideas and suggestions.

## Funding

This work was supported by the Holistique project (défi clé RIVOC Occitanie region, University of Montpellier, France): “*Hyalomma marginatum* in Occitanie region: analysis of biological invasion and associated risks”.

The salary of Charlotte Joly-Kukla, the PhD student working on this project, was funded by INRAE, France.

## Additional Declarations

No competing interests reported.

## Notes

### Competing Interest Statement

The authors have declared no competing interest.

