## Supplementary figures and images for "Contrasting temporal patterns and associations in *Hyalomma marginatum* microbial communities: key insights for the development of novel tick and tick-borne diseases control tools"

### Figure S1

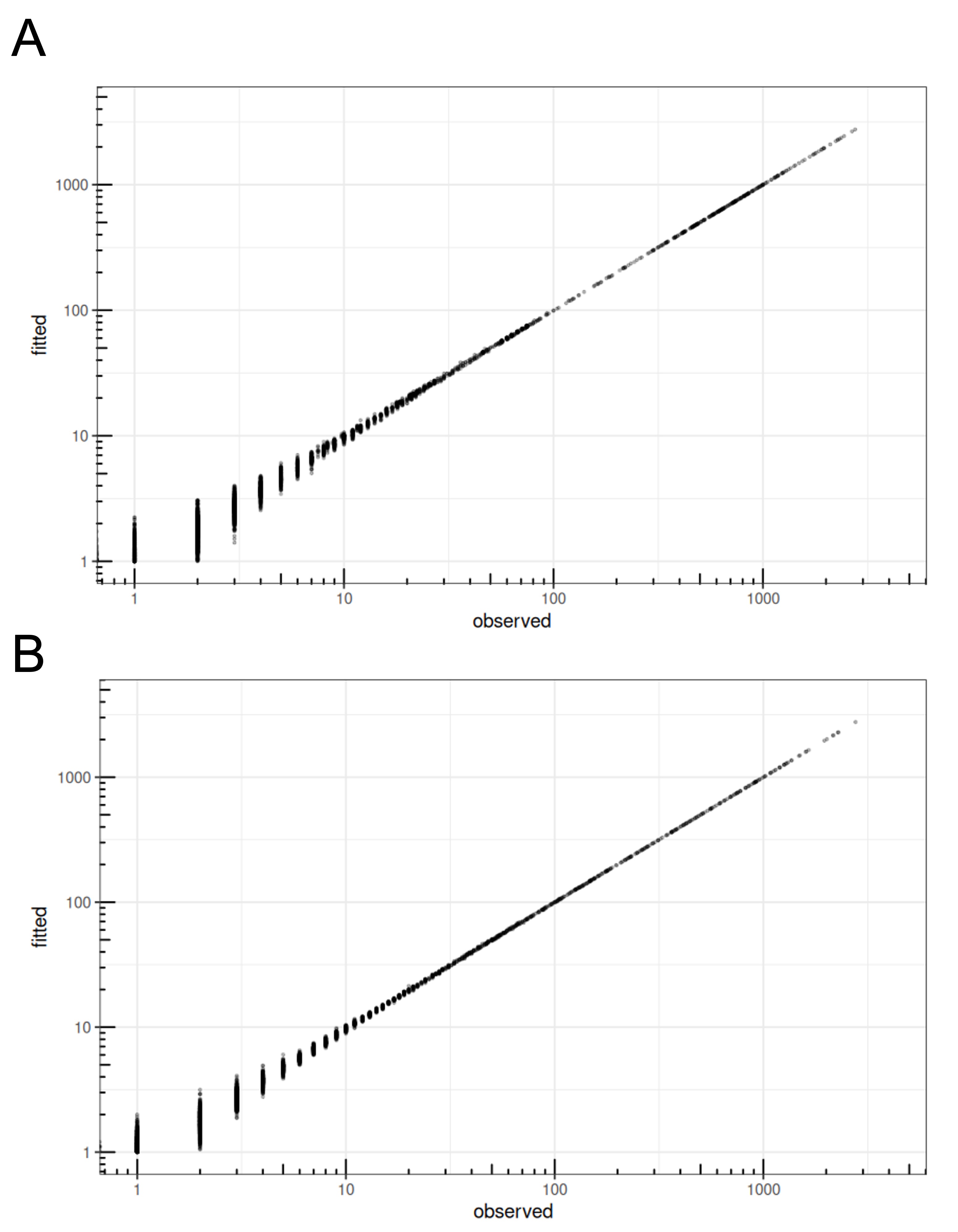

### Figure S2

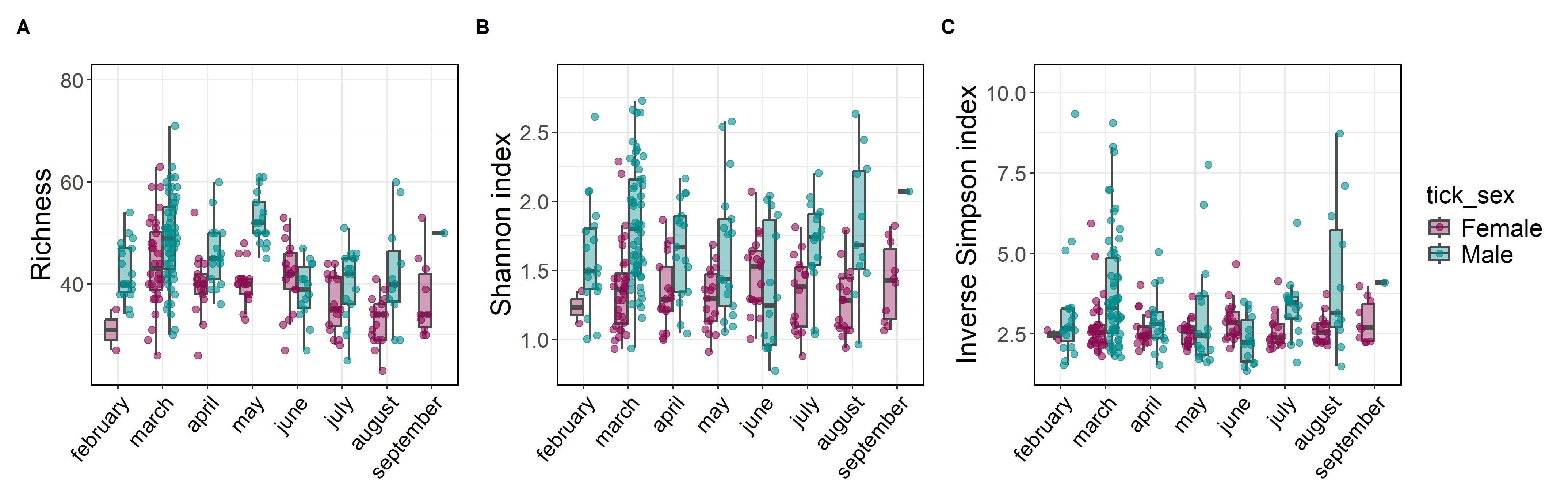

### Figure S3

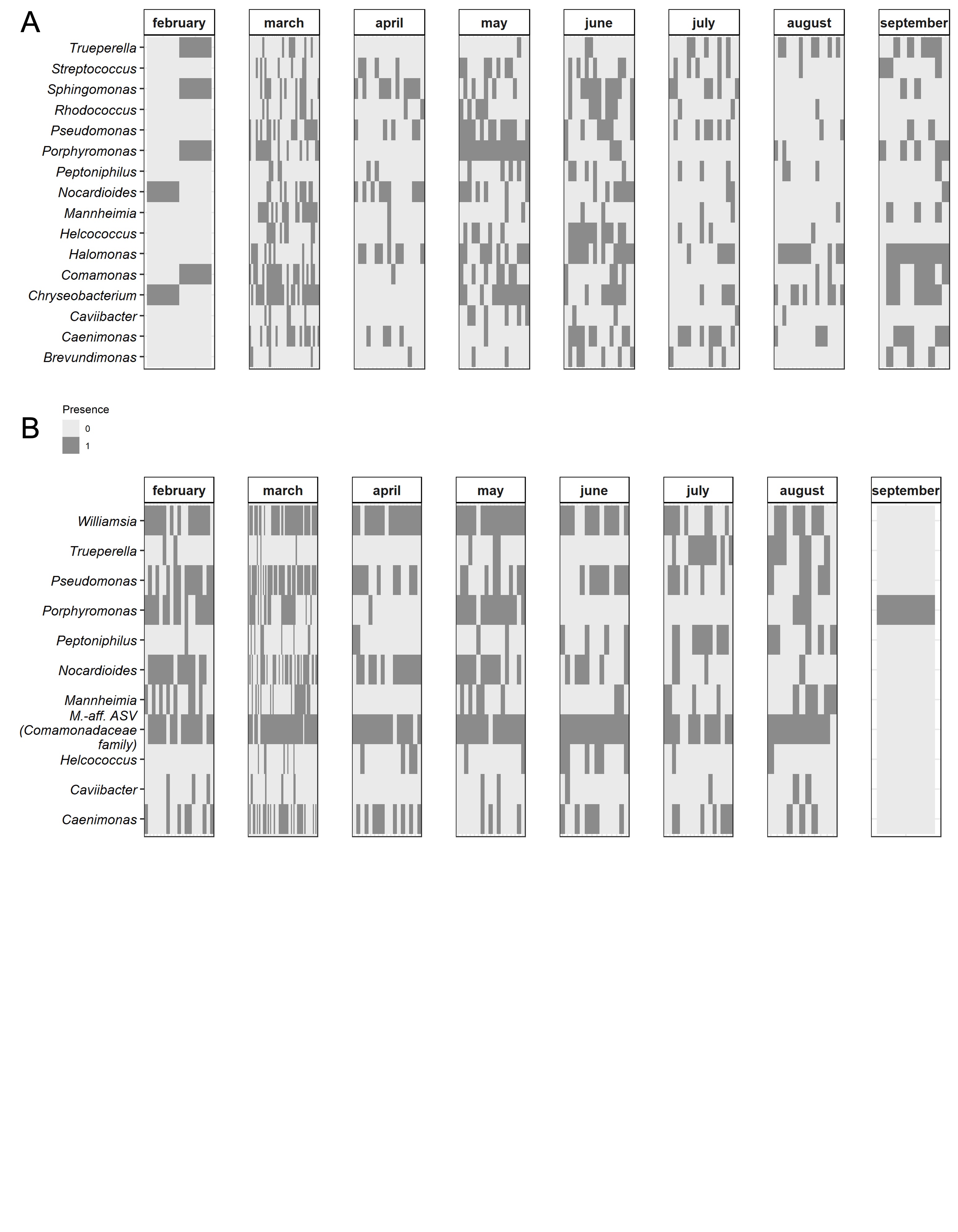

### Figure S4

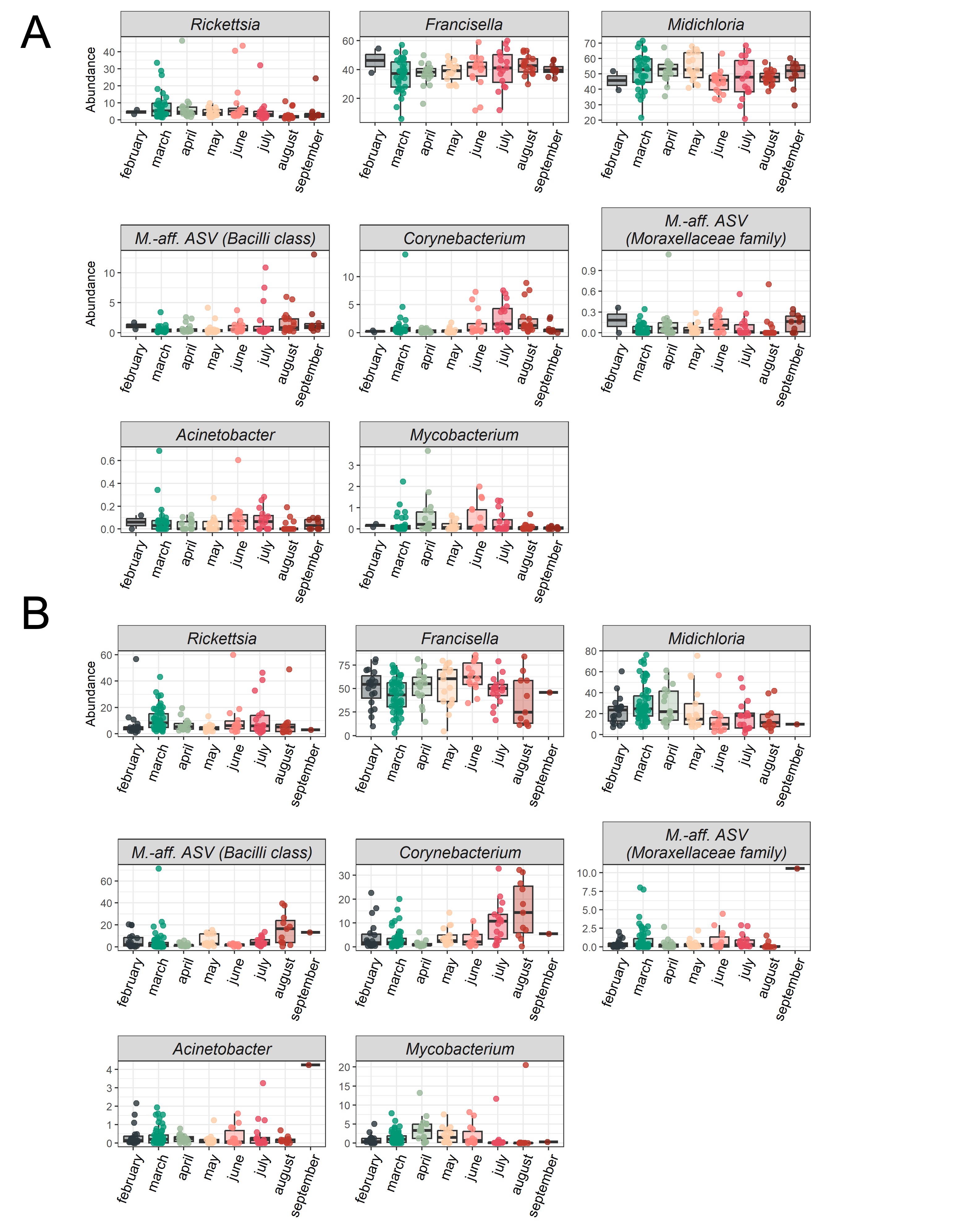

### Figure S5

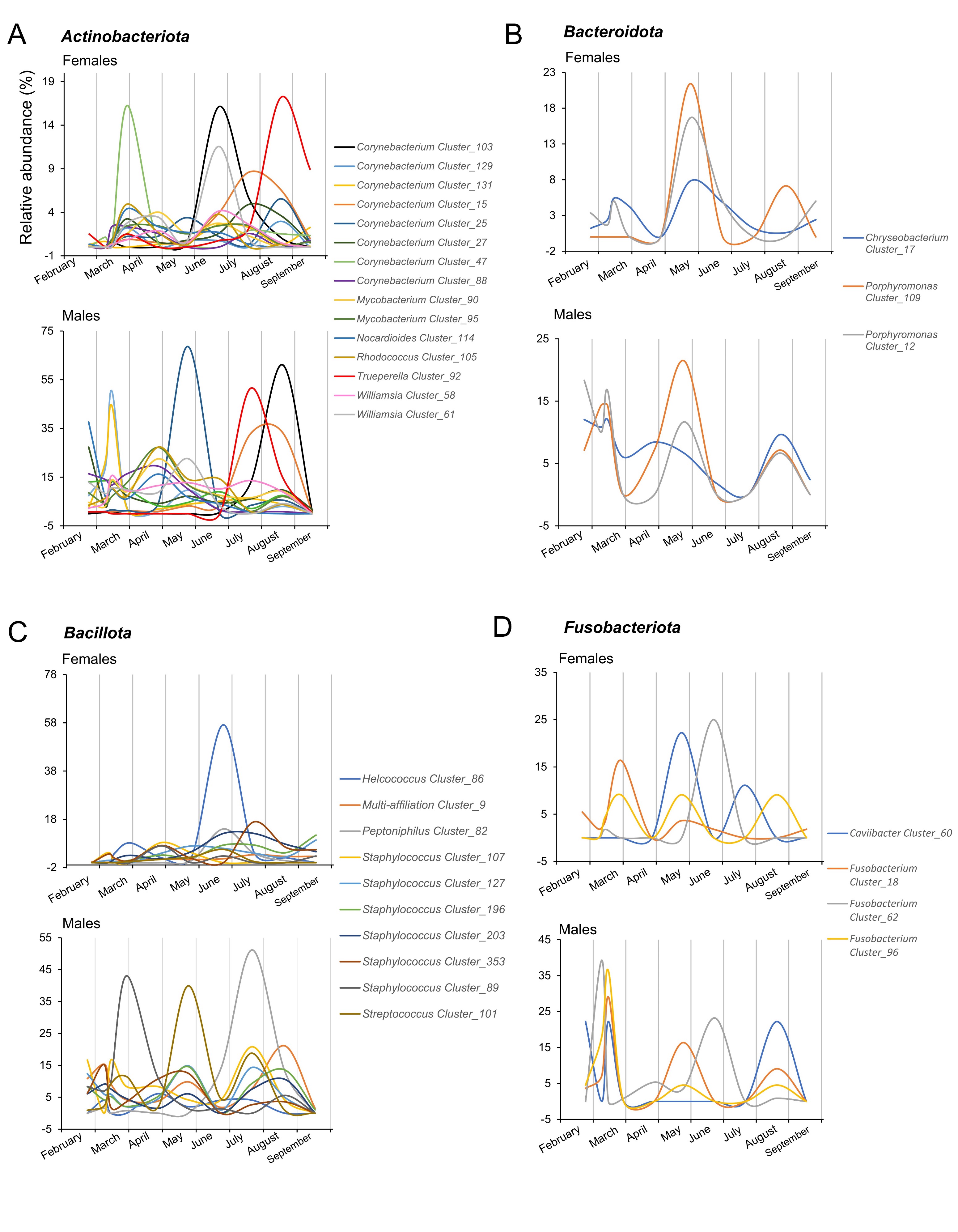

### Figure S6

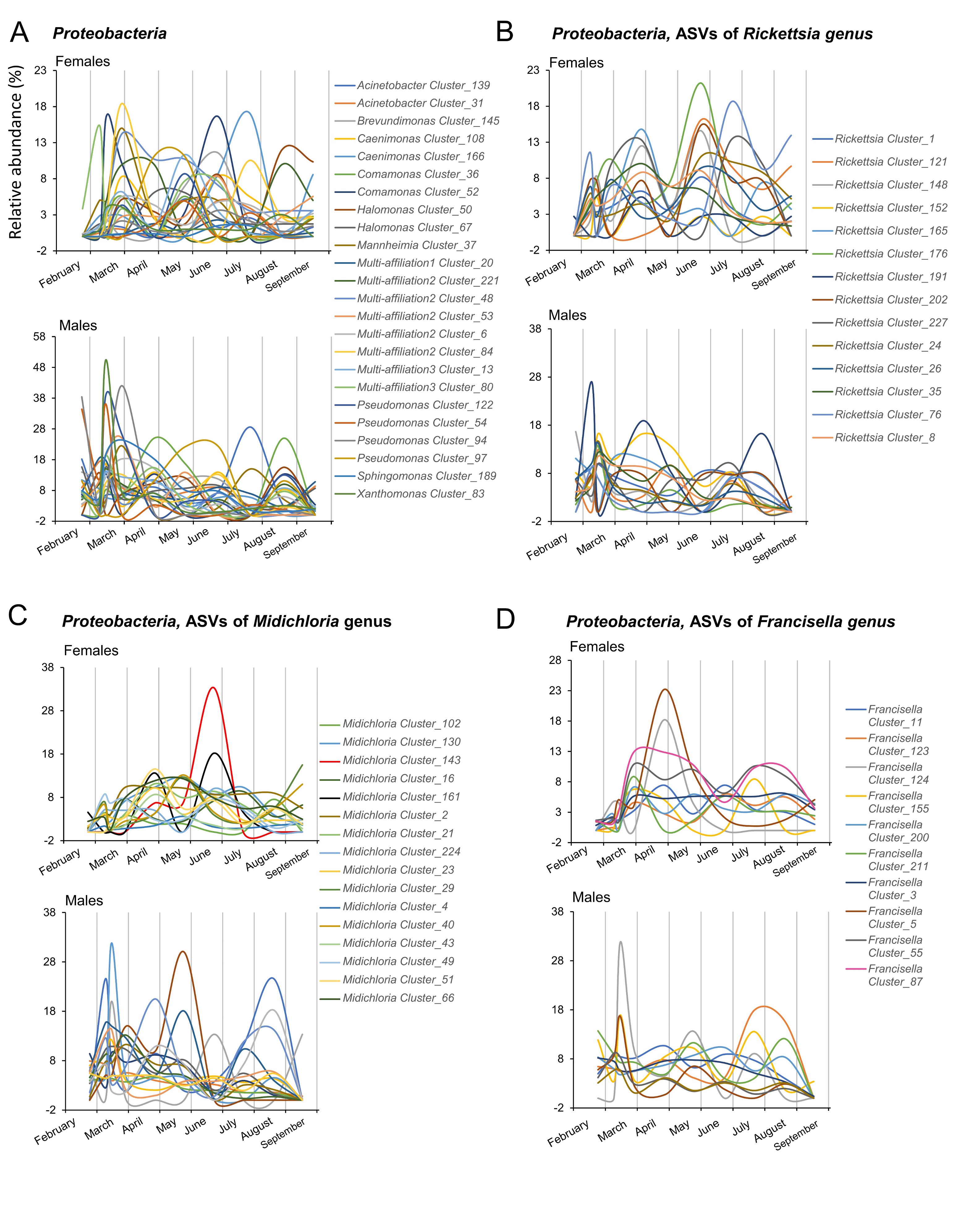

### Figure S7

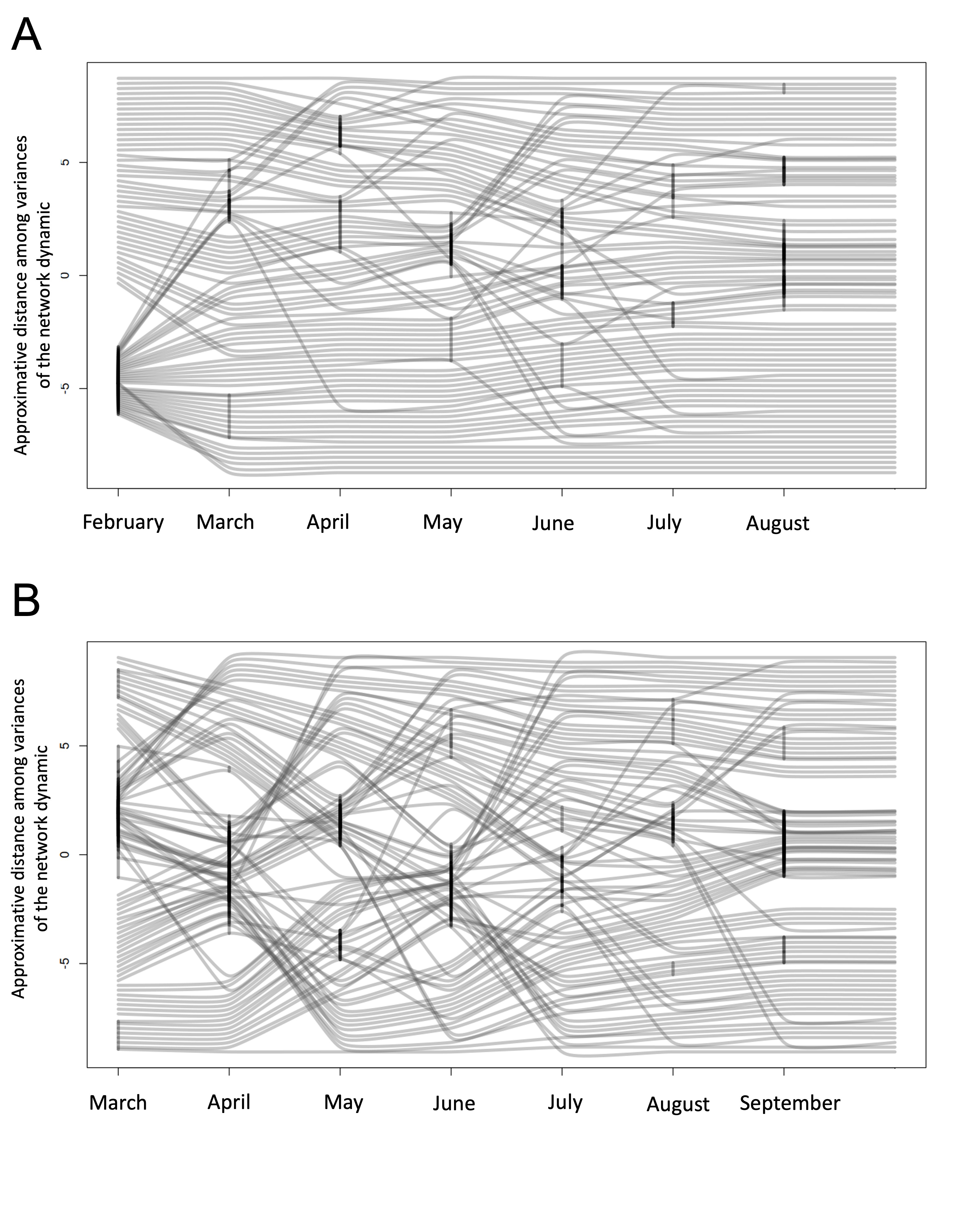
