## Supplementary material for "Contrasting temporal patterns and associations in *Hyalomma marginatum* microbial communities: key insights for the development of novel tick and tick-borne diseases control tools": Table S1

**Supplementary Tables**

**Table S1**: SIMPER analysis (similarity percentage procedure) based on the Bray-Curtis dissimilarities. SIMPER contribution of ASVs to the difference of bacterial communities between males and females is represented in %.

| ASV | Phylum | Genus | p-value | Contribution percentage (%) |
| --- | --- | --- | --- | --- |
| ASV_2 | *Pseudomonadota* | *Midichloria* | 0.001 | 35,0 |
| ASV_3 | *Pseudomonadota* | *Francisella* | 0.001 | 21,8 |
| ASV_189 | *Pseudomonadota* | *Sphingomonas* | 0.003 | 0,8 |
| ASV_55 | *Pseudomonadota* | *Francisella* | 0.001 | 0,4 |
| ASV_31 | *Pseudomonadota* | *Acinetobacter* | 0.030 | 0,3 |
| ASV_13 | *Pseudomonadota* | Multi-affiliated ASV (*Enterobacterales* order) | 0.039 | 0,2 |
| ASV_66 | *Pseudomonadota* | *Midichloria* | 0.001 | 0,2 |
| ASV_87 | *Pseudomonadota* | *Francisella* | 0.001 | 0,2 |
| ASV_130 | *Pseudomonadota* | *Midichloria* | 0.001 | 0,1 |
| ASV_176 | *Pseudomonadota* | *Rickettsia* | 0.010 | 0,1 |
| ASV_121 | *Pseudomonadota* | *Rickettsia* | 0.004 | 0,0 |
| ASV_9 | *Bacillota* | Multi-affiliated ASV (*Bacilli* class) | 0.019 | 5,4 |
| ASV_127 | *Bacillota* | *Staphylococcus* | 0.047 | 0,2 |
| ASV_86 | *Bacillota* | *Helcococcus* | 0.008 | 0,1 |
| ASV_95 | *Actinomycetota* | *Mycobacterium* | 0.004 | 0,4 |
