## Supplementary material for "Contrasting temporal patterns and associations in *Hyalomma marginatum* microbial communities: key insights for the development of novel tick and tick-borne diseases control tools": Table S2

**Supplementary Tables**

**Table S2**: Analysis of the influence of month on the relative abundance of the most abundant genera (<2.5%) in both males and females.

| Genus | Male | Female |
| --- | --- | --- |
| *Francisella* | Yes, *χ*^2^=18.01, p=0.01 | No*, χ*^2^=7.55, p=0.37 |
| *Midichloria* | Yes, *χ*^2^=21.36, p=0.003 | No*, χ*^2^=11.80, p=0.11 |
| *Rickettsia* | No*, χ*^2^=11.20, p=0.13 | No*, χ*^2^=13.09, p=0.06 |
| *Corynebacterium* | Yes, *χ*^2^=55.27, p=1.32 x 10^-9^ | Yes, *χ*^2^=62.99, p=3.81 x 10^-11^ |
| Multi-affiliated ASV (class *Bacilli*) | Yes, *χ*^2^=39.97, p=1.28 x 10^-6^ | Yes, *χ*^2^=39.97, p=1.27 x 10^-6^ |
| Multi-affiliated ASV (family *Moraxellaceae*) | Yes, *χ*^2^=23.53, p=1.38 x 10^-3^ | Yes, *χ*^2^=14.43, p=0.04 |
| *Acinetobacter* | Yes, *χ*^2^=18.54, p=9.75 x 10^-3^ | No, *χ*^2^=7.10, p=0.42 |
| *Mycobacterium* | Yes*, χ*^2^=16.87, p=0.02 | Yes, *χ*^2^=22.85, p=1.80 x 10^-3^ |

*Notes*. Statistical difference Yes/No (statistics, *df* = 7). We were not able to evaluate the influence of the month for *Williamsia* in males due to technical issues.
